# Nucleotide Biosynthesis Links Glutathione Metabolism to Ferroptosis Sensitivity

**DOI:** 10.1101/2021.07.14.452394

**Authors:** Amy Tarangelo, Joon Tae Kim, Jonathan Z. Long, Scott J. Dixon

## Abstract

Nucleotide synthesis is a metabolically demanding process essential for cell division. Several anti-cancer drugs that inhibit nucleotide metabolism induce apoptosis. How inhibition of nucleotide metabolism impacts non-apoptotic cell death is less clear. Here, we report that inhibition of nucleotide metabolism by the p53 pathway is sufficient to suppress the non-apoptotic cell death process of ferroptosis. Mechanistically, stabilization of wild-type p53 and induction of the p53 target gene *CDKN1A* (p21) leads to decreased expression of the ribonucleotide reductase (RNR) subunits *RRM1* and *RRM2*. RNR is the rate-limiting enzyme of de novo nucleotide synthesis that reduces ribonucleotides to deoxyribonucleotides in a glutathione-dependent manner. Direct inhibition of RNR conserves glutathione which can then be used to limit the accumulation of toxic lipid peroxides, preventing the onset of ferroptosis. These results support a mechanism linking p53-dependent regulation of nucleotide metabolism to non-apoptotic cell death.

## Introduction

Proliferating cells must synthesize large quantities of nucleotides to replicate their DNA. De novo nucleotide synthesis in mammalian cells is accomplished through a multi-step pathway whose rate limiting step is the conversion of ribonucleotide triphosphates (NTPs) to deoxyribonucleotide triphosphates (dNTPs) (Elledge *et al*, 1992; Tran *et al*, 2019). The conversion of NTPs to dNTPs is catalyzed by ribonucleotide reductase (RNR), a heterodimer of RRM1 and RRM2 or RRM2B subunits. The reaction catalyzed by RNR requires reducing equivalents supplied by (reduced) glutathione (GSH) or thioredoxin to complete the catalytic cycle, with the glutathione/glutaredoxin system being most important for mammalian RNR reduction and function (Sengupta *et al*, 2019; Zahedi Avval & Holmgren, 2009). RNR subunit expression is controlled by the cell cycle machinery to match the need for nucleotide synthesis to cell proliferation (Eriksson *et al*, 1984; Sigoillot *et al*, 2003). Thus, cell cycle progression requires redox-regulated RNR function. Whether disruption of cell cycle progression alters other redox-dependent processes in the cell is poorly understood.

The p53 tumor suppressor pathway is an important regulator of cell cycle progression and nucleotide metabolism (Huang *et al*, 2018; Sherley, 1991; Zhang *et al*, 2001). Activation of the p53 pathway may limit oxidative stress and promote cell survival during serine deprivation, possibly by modulating nucleotide synthesis (Maddocks *et al*, 2013). However, the specific mechanism involved in this protective effect is unclear. More generally, how p53 pathway-dependent regulation of nucleotide metabolism links to the broader control of cell fate by p53 is poorly understood.

In addition to its well-established role in apoptosis, the p53 pathway regulates ferroptosis sensitivity (Chu *et al*, 2019; Jiang *et al*, 2015; Kruiswijk *et al*, 2015; Tarangelo *et al*, 2018; Venkatesh *et al*, 2020a; Xie *et al*, 2017). Ferroptosis is an oxidative, iron-dependent form of cell death important for tumor suppression and pathological cell death in mammals (Badgley *et al*, 2020; Fang *et al*, 2019; Linkermann *et al*, 2014; Skouta *et al*, 2014; Ubellacker *et al*, 2020). Ferroptosis is characterized by the accumulation of toxic lipid peroxides at the cell membrane (Dixon *et al*, 2012; Stockwell *et al*, 2017). Lipid peroxide accumulation is inhibited by glutathione peroxidase 4 (GPX4), a lipid hydroperoxidase that uses reduced glutathione (GSH) as a cofactor (Friedmann Angeli *et al*, 2014; Yang *et al*, 2014). Ferroptosis can therefore be induced by depriving cells of the sulfur-containing GSH precursor, cysteine, via inhibition of the system x_c_^-^ cystine/glutamate antiporter (e.g. using small molecules of the erastin family), or by direct inhibition of GPX4 itself (Forcina & Dixon, 2019). Stabilization of wild-type p53 and induction of p21 can decrease the susceptibility of cancer cells to ferroptosis by conserving intracellular glutathione (Tarangelo *et al*., 2018). Overexpression of p21 alone may also be sufficient to inhibit ferroptosis via interaction with cyclin dependent kinases (CDKs) in some contexts (Venkatesh *et al*, 2020b). However, the specific mechanism by which p53 and p21 conserve glutathione to inhibit ferroptosis is not clear.

Here, we explored how stabilization of p53 and induction of p21 promote ferroptosis resistance in human cancer cells. Using transcriptomic analysis, we find that activation of the p53-p21 pathway downregulates expression of many genes involved in nucleotide metabolism. Among the most downregulated genes in this subset are the RNR subunits *RRM1* and *RRM2*. Using cell death kinetic analysis, gene-editing and metabolic tracing, we show that inhibition of RNR-dependent nucleotide metabolism allows for conservation of intracellular GSH and a reduction in ferroptosis susceptibility in cancer cells. Thus, regulation of nucleotide synthesis by the p53-p21 axis provides a crucial link between GSH metabolism and ferroptosis susceptibility.

## Results

### Pharmacological p53 stabilization inhibits ferroptosis

The effects of p53 on ferroptosis are controversial, with evidence that p53 may induce or suppress this cell death process in a context-dependent manner (Jiang *et al*., 2015; Tarangelo *et al*., 2018; Xie *et al*., 2017). We examined the effects of p53 manipulation in human HT-1080 fibrosarcoma cells, which express wild-type p53. We inhibited the function of the p53 negative regulator human double minute 2 (HDM2) using the potent and specific small molecule inhibitors nutlin-3 and MI-773/SAR405838 (Vassilev *et al*, 2004; Wang *et al*, 2014). In all experiments cell death was quantified using the scalable time-lapse analysis of cell death kinetics (STACK) imaging method (Forcina *et al*, 2017). Using STACK, we determined the precise time of cell death onset within each population, denoted as D_O_. Consistent with previous results (Tarangelo *et al*., 2018), nutlin-3 and MI-773 stabilized p53 levels and delayed the onset of ferroptosis in response to erastin2 treatment in p53 wild-type HT-1080 cells, while an inactive enantiomer of nutlin-3 (nutlin-3b) had no such effects (**Figure 1A-C**). Cell death in these experiments was suppressed by co-treatment with the specific ferroptosis inhibitor ferrostatin-1, confirming that p53 stabilization did not alter the mode of cell death induced by erastin2 (**Figure 1C**). The effects of nutlin-3 were dose-dependent, with increasing concentrations of this inhibitor resulting in greater p53 stabilization, which correlated with increased induction of the p53 target genes *CDKN1A* and *HDM2* and greater suppression of erastin2-induced ferroptosis (**Figure S1A-C**). We also observed increased D_O_ in cells treated with nutlin-3 or MI-773, compared with cells treated with DMSO or nutlin-3b, indicating that cell death was delayed (**Figure 1D, S1D**). Collectively, these results show that pharmacological HDM2 inhibition robustly delays the onset of ferroptosis in HT-1080 cells through an on-target mechanism.

**Figure 1:**
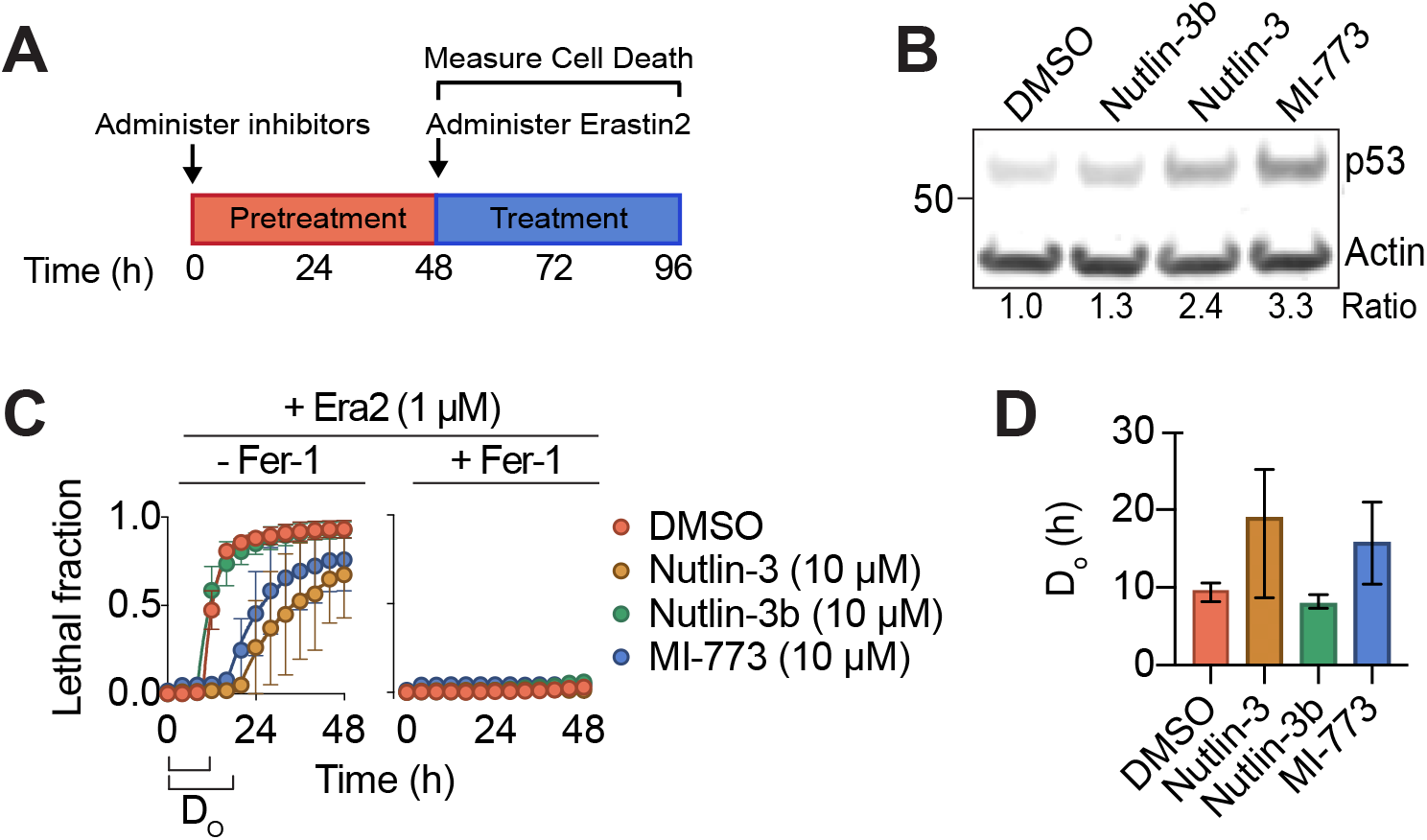
Diverse p53-activating stimuli suppress ferroptosis. **(A)** Schematic of experimental procedures for assaying cell death susceptibility. **(B)** Western blot for HT-1080 Control cells treated with MDM2 inhibitors (10 μM, 48 h). **(C)** Lethal response curves for HT-1080 Control cells expressing a nuclear mKate2 signal (HT-1080^N^) pretreated with MDM2 inhibitors (10 μM, 48 h) followed by treatment with Erastin2 (Era2, 1 μM), ± the specific ferroptosis suppressor ferrostatin-1 (Fer-1, 1 μM). **(D)** Quantification of the time of death onset (D_O_) for the lethality curves shown in **C**, calculated using STACK. All data are plotted as the mean ± SD of ≥3 independent experiments. D_O_ values are calculated using the STACK approach and plotted as a best-fit value ± 95% confidence interval (CI).

### p21 is necessary but not sufficient to suppress ferroptosis

We previously showed that expression of *CDKN1A*, encoding the cyclin dependent kinase inhibitor p21, was necessary for wild-type p53 to suppress ferroptosis (Tarangelo *et al*., 2018). Recently it was suggested that p21 expression alone may be sufficient to suppress ferroptosis in some contexts, independent of p53 (Venkatesh *et al*., 2020b). To examine whether p21 expression was sufficient to fully recapitulate the functions of p53 in ferroptosis suppression, we generated cell lines expressing Flag-tagged *CDKN1A* under the control of a doxycycline (Dox)-inducible promoter, denoted p21^Dox^. We introduced p21^Dox^ into both HT-1080 Control (i.e. wild-type) and p53 knockout (p53^KO^) backgrounds (Tarangelo *et al*., 2018). Dox treatment induced robust expression of full-length p21 in both Control and p53^KO^ cells, which was not further augmented by co-treatment with nutlin-3 (**Figure 2A**). Dox pretreatment alone delayed the onset of ferroptosis (i.e. increased D_O_) in response to erastin2 in Control cells capable of expressing p21, albeit not to the same extent as nutlin-3 treatment (**Figure 2B,C).** By contrast, in p53^KO^ cells, Dox-induced p21 induction had little ability to prevent ferroptosis (**Figure 2B,C)**. Thus, p21 expression alone is not sufficient to fully recapitulate the effects of p53 stabilization, especially when cells lack wild-type p53.

**Figure 2:**
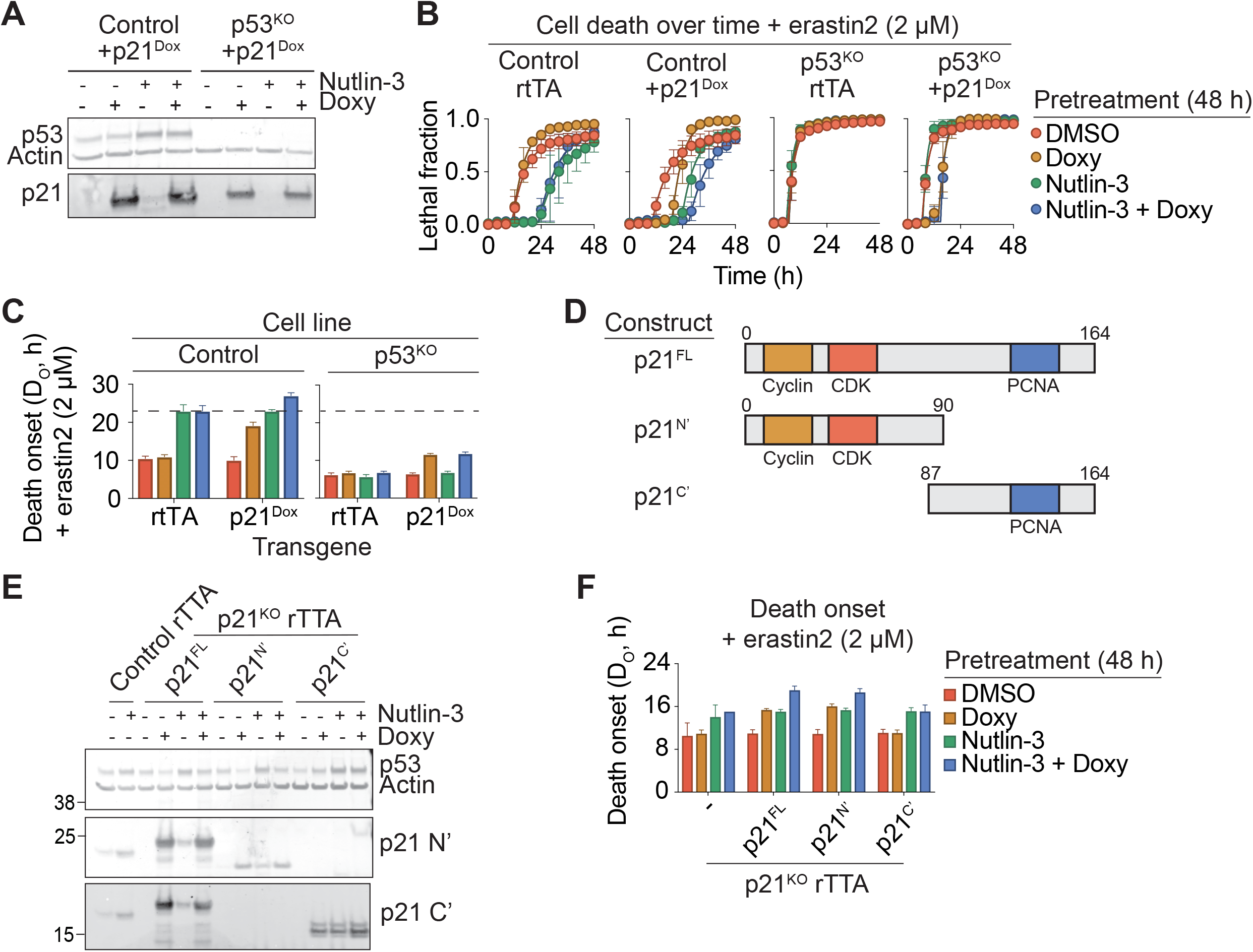
The CDK-binding domain of p21 is required to suppress ferroptosis. **(A)** Protein expression in HT-1080 Control or p53^KO^ + p21^Dox^ treated ± nutlin-3 (10 μM) ± doxycycline (Doxy, 1 μg/mL) for 48 h. **(B)** Cell death in HT-1080^N^ Control or p53^KO^ cells ± the p21^Dox^ cassette, pretreated ± nutlin-3 (10 μM) ± doxycycline (1 μg/mL) for 48 h, followed by treatment with Era2 (2 μM). Data represent mean ± SD. **(C)** Timing of death onset (D_O_) calculated from data shown in **B**. **(D)** Schematic of the structure of p21 full length (FL), N-terminus (N’) or C-terminus (C’). **(E)** Protein expression in HT-1080 stable cell lines ± nutlin-3 (10 μM) ± doxycycline (1 μg/mL) for 48 h. **(F)** Cell death onset (D_O_) plotted for the indicated conditions. All results are from three independent experiments. Data in **C** and **F** represent mean ± 95% confidence interval (CI).

We further dissected the contribution of different p21 functional domains to ferroptosis suppression. In particular, we focused on the p21 N-terminal cyclin-CDK binding domain, required for inhibition of G1/S cell cycle progression, and the C-terminal PCNA binding domain involved in suppressing DNA synthesis (Chen *et al*, 1995) (**Figure 2D**). We introduced Dox-inducible full-length (FL) Flag-tagged p21, or isolated N and C terminal truncation mutants (p21^N’^ and p21^C’^, respectively), into HT-1080 p21 knockout (KO) cells (Tarangelo *et al*., 2018). Dox induction of all constructs was validated with antibodies specific for epitopes in the p21 N and C termini (**Figure 2E).** Consistent with negative feedback by p21 on p53 levels (Broude *et al*, 2007), induction of p21^FL^ and p21^N’^, but not p21^C’^, decreased basal p53 levels and attenuated p53 stabilization upon nutlin-3 treatment, further validating the function of these proteins (**Figure 2E)**. With respect to cell death, induction of p21^FL^ modestly delayed the onset of erastin2-induced death alone, and to a greater extent when combined with nutlin-3, consistent with an independent requirement for both p53 and p21 in the suppression of ferroptosis (**Figure 2F)**. Induction of p21^N’^, but not p21^C’^, likewise modestly suppressed ferroptosis alone and more potently suppressed ferroptosis together with nutlin-3 treatment, to the same extent as p21^FL^. Thus, the p21 N-terminus is required to suppress ferroptosis.

### p53 does not modulate ferroptosis via glutathione export or altered electron transport chain function

Activation of the p53-p21 pathway can suppress ferroptosis by conserving intracellular glutathione (Tarangelo *et al*., 2018). The specific mechanism accounting for glutathione conservation is unclear. To investigate further, we initially pursued a hypothesis-driven approach. First, we investigated the role of *ABCC1*, which encodes the multidrug-resistance protein 1 (MRP1). MRP1 exports glutathione from the cell and loss of MRP1 expression can thereby conserve intracellular glutathione and delay the onset of ferroptosis, much like p53 stabilization (Cao *et al*, 2019; Cole, 2014). Accordingly, we investigated whether p53 stabilization delayed ferroptosis and conserved intracellular glutathione by downregulating *ABCC1* expression. However, nutlin-3 treatment did not decrease *ABCC1* expression in HT-1080 Control cells or in p53 wild-type U-2 OS osteosarcoma cells (**Figure S2A**). Indeed, while nutlin-3 pretreatment suppressed erastin2-induced cell death in U-2 OS cells expressing an empty control vector, this condition enhanced ferroptosis in U-2 OS cells overexpressing MRP1 (**Figure S2B**). Thus, p53 did not appear to suppress ferroptosis by reducing *ABBC1*/MRP1 function.

Mitochondria are a major source of intracellular ROS, whose accumulation is buffered by glutathione-dependent enzymes (Handy *et al*, 2009; Mari *et al*, 2009). Mitochondrial oxidative phosphorylation has been suggested to promote ferroptosis through the production of lipid peroxides (Gao *et al*, 2019). p53 activation can suppress the mitochondrial tricarboxylic acid (TCA) cycle and oxidative phosphorylation through inhibition of malic enzymes 1/2 or by inducing a PUMA-dependent metabolic switch (Jiang *et al*, 2013; Kim *et al*, 2019). Accordingly, we hypothesized that p53 stabilization conserved glutathione and suppressed ferroptosis by reducing mitochondrial activity. To investigate this, we generated HT-1080 cells lacking mitochondrial DNA (i.e. ρ^0^ cells) via long-term incubation in ethidium bromide. Compared with control (i.e. ρ^+^) cells, ρ^0^ cells expressed mitochondrial genes at undetectable levels and were entirely deficient in oxidative metabolism, as detected using Seahorse technology (**Figure S2C,D**). However, there was no difference in basal sensitivity to erastin2-induced ferroptosis in ρ^0^ compared to ρ^+^ cells, and pretreatment with nutlin-3 suppressed ferroptosis equally well in both lines (**Figure S2E**). These results indicated that the ability of the p53 pathway to suppress ferroptosis was unlikely to involve changes in mitochondrial oxidative phosphorylation.

### The p53-p21 axis regulates nucleotide metabolic gene networks

We next took an unbiased approach to identify candidate pathways impacting glutathione metabolism and ferroptosis sensitivity downstream of p53 and p21. Towards this end, we performed RNA sequencing of HT-1080 Control, p53^KO^ and p21^KO^ treated with or without nutlin-3 (10 μM, 48 h). In Control cells, nutlin-3 treatment resulted in downregulation of 2,714 genes and upregulation of 2,429 genes (**Figure 3A**). Consistent with expectations, bona fide p53 targets including *HDM2* and *CDKN1A* were among the most highly upregulated genes in Control cells, while genes known to be negatively regulated by p53, such as *E2F2*, were strongly downregulated (Chen *et al*, 1994; el-Deiry *et al*, 1993; Timmers *et al*, 2007). Compared to Control cells, p53^KO^ cells exhibited far fewer nutlin-3-induced changes in gene expression, consistent with most changes observed in Control cells being p53-dependent (**Figure 3A**). By contrast, in p21^KO^ cells, nutlin-3 treatment resulted in the differential expression of even more genes than observed in Control cells, consistent with the notion that p21 is itself an important transcriptional regulator (**Figure 3A**) (Delavaine & La Thangue, 1999; Ferrandiz *et al*, 2012). We exploited this large RNA Seq dataset to develop specific hypotheses concerning the regulation of ferroptosis by the p53 pathway.

**Figure 3:**
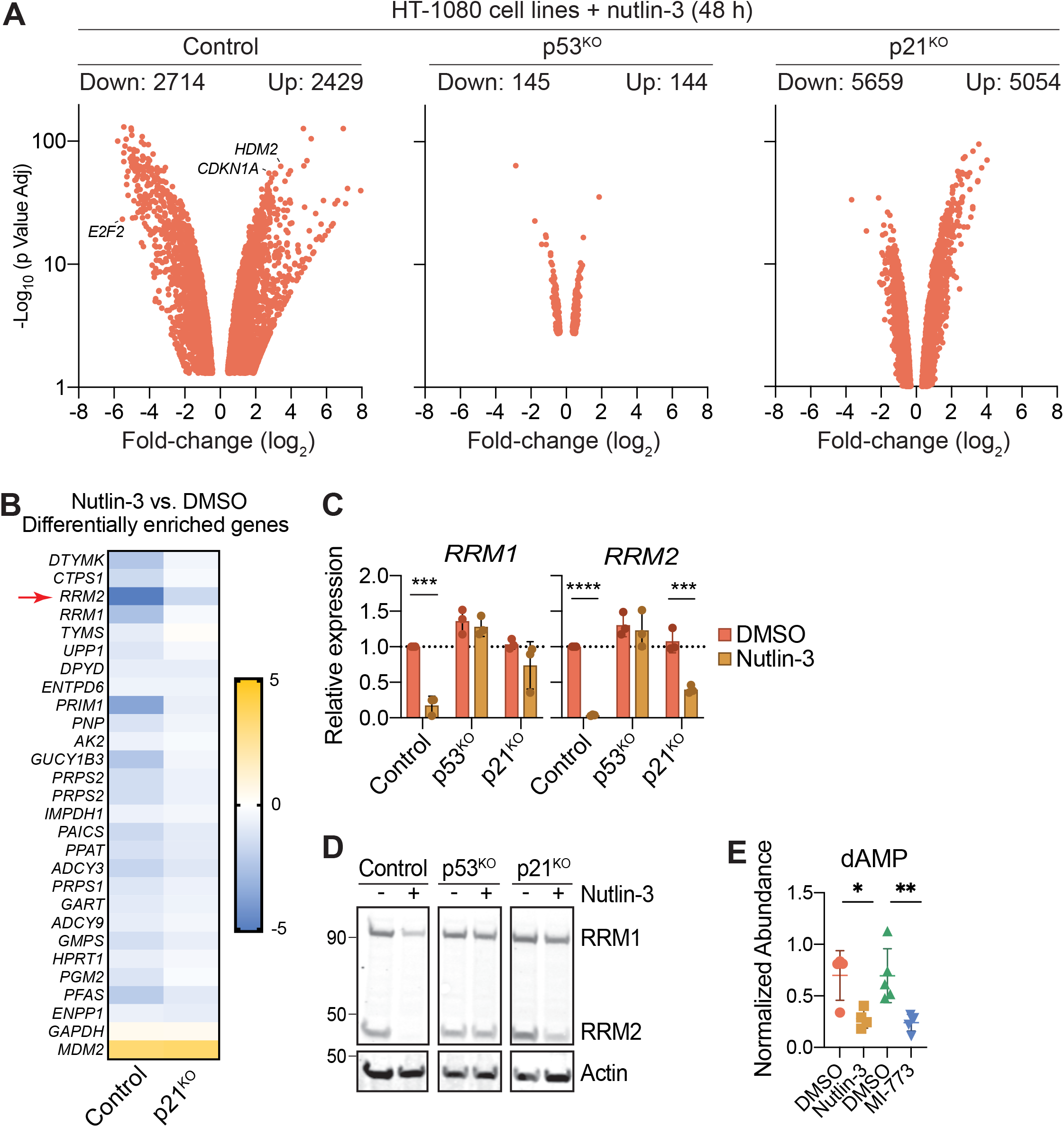
p53 stabilization reduces RNR expression and activity. **(A)** Volcano plot summarizing significantly (*P* < 0.05) upregulated and downregulated genes in HT-1080 Control, p53^KO^ and p21^KO^ cells treated ± nutlin-3 (10 μM, 48 h). RNA sequencing data is the average of two independent experiments. **(B)** Differential gene expression summary for genes with annotated functions in nucleotide metabolism from cells treated and analyzes as described in **A**. **(C)** mRNA levels in HT-1080 Control, p53^KO^, and p21^KO^ cells treated ± nutlin-3 (10 μM, 48 h). **(D)** Protein expression in HT-1080 Control, p53^KO^, and p21^KO^ cells treated ± nutlin-3 (10 μM, 48 h). **(E)** Levels of dAMP detected by LC-MS in HT-1080 Control cells treated with DMSO, nutlin-3 (10 μM) or MI-773 (10 μM) for 48 h. Data in **C** and **E** are mean ± SD of ≥3 biological replicates.

In Control cells, nutlin-3 treatment downregulated numerous genes involved in nucleotide metabolism, including *DTYMK*, *CTPS1*, *RRM1* and *RRM2* (**Figure 3B**). *RRM1* and *RRM2* encode the R1 and R2 subunits of the heterodimeric enzyme ribonucleotide reductase (RNR). RNR catalyzes the reduction of ribonucleotides to deoxyribonucleotides. In mammalian cells, RNR activity requires GSH (Sengupta *et al*., 2019; Zahedi Avval & Holmgren, 2009). Activation of the p53/p21 pathway can enable cancer cells to resist oxidative stress, and this effect may involve inhibition of nucleotide synthesis (Maddocks *et al*., 2013). We therefore focused on the potential regulation of RNR function by the p53 pathway. Using RT-qPCR we confirmed that both *RRM1* and *RRM2* were downregulated following nutlin-3 treatment, with *RRM2* downregulation being entirely p53-dependent and *RRM1* regulated by both p53 and p21 (**Figure 3C**). At the protein level, R2 expression was more sensitive than expression of R1 to downregulation upon p53 stabilization (**Figure 3D**). Consistent with decreased RNR activity, treatment of Control cells with nutlin-3 or MI-773 decreased levels of dAMP, a metabolite downstream of RNR (**Figure 3E**). Thus, activation of the p53-p21 pathway broadly downregulates nucleotide metabolism, including RNR expression and activity.

### RNR inhibition blocks ferroptosis

RNR is oxidized as part of its catalytic mechanism. In mammalian cells, reduction of oxidized RNR requires glutathione (Sengupta *et al*., 2019). Accordingly, we hypothesized that inhibition of RNR function may suppress ferroptosis by reducing the draw on the existing glutathione pool, which can then be re-routed to suppress ferroptosis. To test this hypothesis, we first treated cells with small molecule inhibitors of pyrimidine and purine metabolism, which generate the ribonucleotide precursors that RNR reduces to dNTPs in a GSH-dependent manner. Cells were pretreated with mycophenolic acid (MPA), to inhibit the purine synthetic enzyme inosine-5’-monophosphate dehydrogenase (IMPDH), or with pyrazofurin (Pyr), to inhibit the pyrimidine synthetic enzyme orotidine 5’-monophosphate decarboxylase (UMPS) (Fleming *et al*, 1996; Ringer *et al*, 1991). Nucleotide depletion itself triggers p53 stabilization, and we confirmed that both MPA and Pyr caused this effect in HT-1080 Control cells (**Figure 4A**). Consistent with our hypothesis, 24 h pretreatment with either MPA or Pyr subsequently protected Control cells from erastin2-induced cell death (**Figure 4B**). Moreover, unlike with nutlin-3 (**Figure 1C**) (Tarangelo *et al*., 2018), MPA and Pyr pretreatment protected both p53^KO^ and p21^KO^ cells from subsequent erastin2 treatment, consistent with RNR acting downstream of the p53/p21 axis. Crucially, in all three cell lines, the de novo GSh synthesis inhibitor buthionine sulfoximine (BSO) abolished the ability of MPA and Pyr pretreatment to inhibit ferroptosis, consistent with a GSH-dependent protective mechanism (**Figure 4B**). Of note, the restoration of cell death by BSO co-treatment also indicates that MPA and Pyr were not inhibiting ferroptosis in an off-target manner by acting as radical trapping antioxidants (Conlon *et al*, 2021).

**Figure 4:**
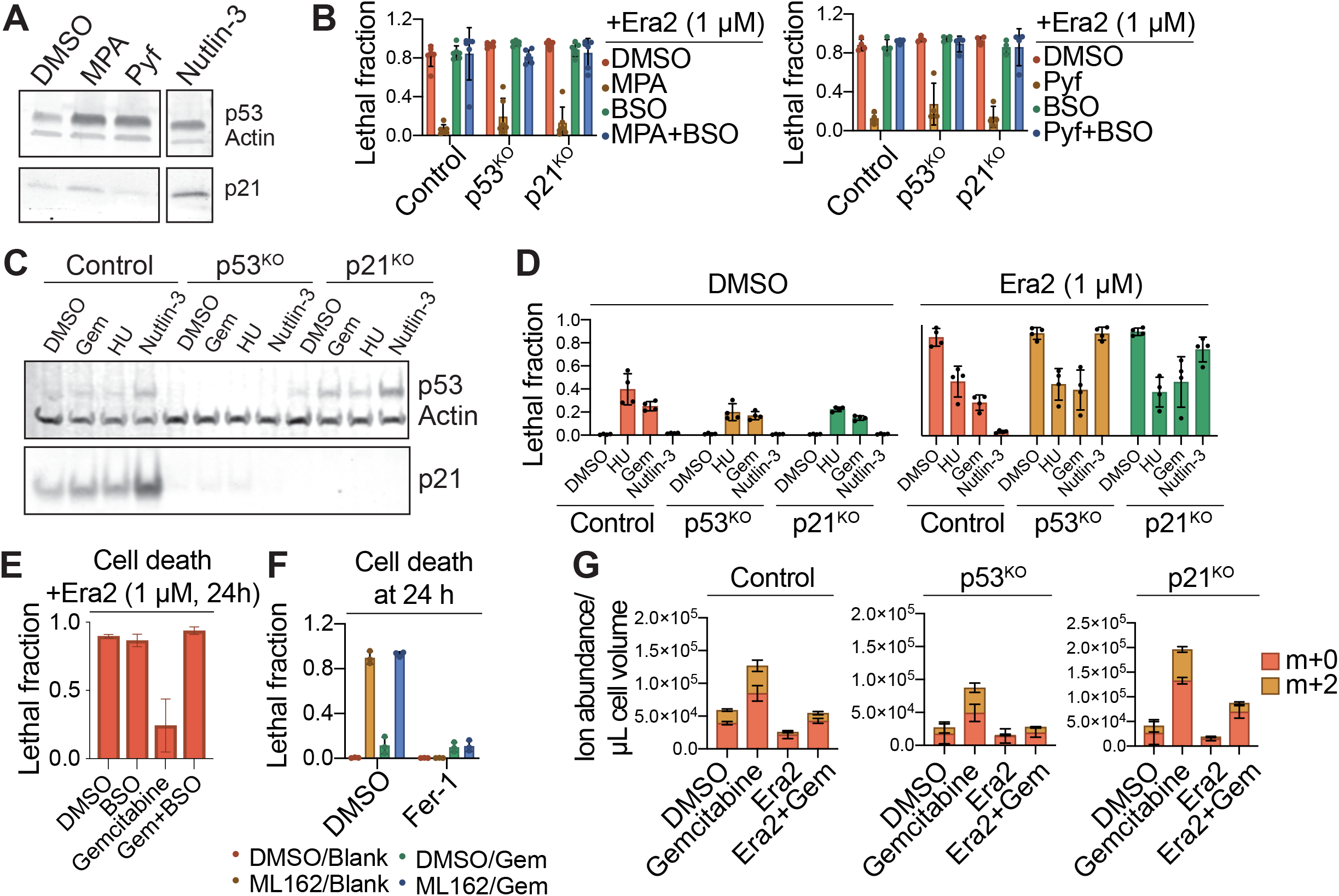
Pharmacological inhibition of nucleotide metabolism modulates ferroptosis. **(A)** Protein expression in HT-1080 Control cells treated with MPA (5 μM), Pyr (5 μM) or Nutlin-3 (10 μM). **(B)** Cell death in HT-1080^N^ Control, p53^KO^ and p21^KO^ cells pretreated for 48 h with mycophenolic acid (MPA, 5 μM, left) or pyrazofurin (Pyr, 5 μM, right) ± buthionine sulfoximine (BSO, 100 μM), then treated with erastin2 (Era2, 1 μM, 24 h). **(C)** Protein expression in HT-1080 Control cells treated with DMSO, hydroxyurea (5 mM), gemcitabine (200 nM) or nutlin-3 (10 μM). **(C)** Cell death in HT-1080^N^ Control, p53^KO^ or p21^KO^ cells pretreated with hydroxyurea (5 mM), gemcitabine (200 nM) or nutlin-3 (10 μM), then ± era2 for 24 h. **(E)** Cell death in HT-1080 Control cells pretreated with gemcitabine (Gem, 200 nM) ± BSO (100 μM), then treated with Era2 (1 μM, 24 h). **(F)** Cell death in HT-1080 Control cells pretreated with ± gemcitabine (200 nM) for 48 h then treated ± ML162 (1 μM) ± ferrostatin-1 (Fer-1, 1 μM). **(G)** Metabolic flux assay using LC-MS detection of GSH in HT-1080 Control, p53^KO^ or p21^KO^ cells pretreated with DMSO or gemcitabine (200 nM) for 48 h. Following pretreatment, media containing Era2 (1 μM) and U-^13^C Serine was added to cells for 8 h. De novo synthesized GSH is labeled as an m+2 isotopologue (yellow). Results in **B,D-G** are mean ± SD of ≥3 independent experiments.

RNR can be directly inhibited using the small molecules gemcitabine (Gem) and hydroxyurea (HU), which serve as analogs of cytidine to inhibit dNTP synthesis (Heinemann *et al*, 1990; Krakoff *et al*, 1968). Gem and HU did not strongly activate p53 and/or induce p21 expression in Control, p53^KO^ or p21^KO^ cells (**Figure 4C**). However, pretreatment with Gem or HU strongly suppressed ferroptosis in all three cell lines (**Figure 4D**). The effect of Gem pretreatment on ferroptosis was reversed by co-treatment with BSO, consistent with a GSH-dependent mechanism (**Figure 4E**). Further consistent with this mechanism, Gem pretreatment did not suppress ferroptosis induced by the direct GPX4 inhibitor ML162, whose ability to induce ferroptosis is independent of intracellular GSH pools (Yang *et al*., 2014) (**Figure 4F**).

Our results suggested that RNR inhibition suppressed ferroptosis by conserving intracellular glutathione. To directly test this model, we assayed GSH synthesis and levels by performing metabolic tracing with uniformly-labelled ^13^C-serine. Serine can be converted to glycine within the cell, and glycine then incorporated into GSH. Thus, GSH synthesized de novo using ^13^C-glycine will be labeled with two heavy carbons and detected as an m+2 isotopologue. Cells were pretreated with Gem for 48 h, then treated with Era2 in medium containing ^13^C-serine. Pretreatment with Gem increased both the total levels and de novo synthesis of GSH in Control, p53^KO^ and p21^KO^ cell lines (**Figure 4G**). Control and p21^KO^ cells pretreated with Gem also conserved a larger portion of the existing (m+0) intracellular GSH pool following treatment with erastin2 compared to cells pretreated with DMSO, with weaker effects observed in p53^KO^ cells (**Figure 4G**). These data suggest that RNR inhibition may suppress ferroptosis by conserving intracellular GSH.

### RNR inhibition is sufficient to suppress lipid ROS accumulation and ferroptosis

To strengthen the results obtained above using pharmacological inhibitors we silenced expression of the essential RNR subunit *RRM1* using short interfering RNA (siRNA). Silencing of *RRM1* in HT-1080 Control cells strongly decreased the corresponding protein levels and led to compensatory upregulation of RRM2, indicating that our silencing reagent worked as expected (**Figure 5A**). Consistent with our pharmacological results, si-*RRM1* treatment resulted in greater conservation of intracellular GSH m+0 pools following erastin2 treatment compared to si-Control-treated cells, as determined using ^13^C-serine metabolic tracing (**Figure 5B**). Silencing of *RRM1* likewise inhibited the accumulation of lipid peroxides in response to erastin2 treatment, as determined using confocal imaging of C11 BODIPY 581/591 (**Figure 5C**). Concurrently, *RRM1* silencing was sufficient to suppress ferroptotic cell death in Control cells treated with erastin2 (**Figure 5D**). This effect of *RRM1* silencing on erastin2-induced ferroptosis was specific, as *RRM1* silencing did not inhibit cell death in response to several mechanistically distinct inducers of apoptosis (camptothecin, bortezomib, staurosporine) or other forms of non-apoptotic cell death (CIL56) that are not directly linked to intracellular glutathione metabolism (Ko *et al*, 2019) (**Figure S3**). Thus, inhibition of RNR function is sufficient to suppress ferroptosis specifically in response to system x_c_^-^ inhibition in Control cells.

**Figure 5:**
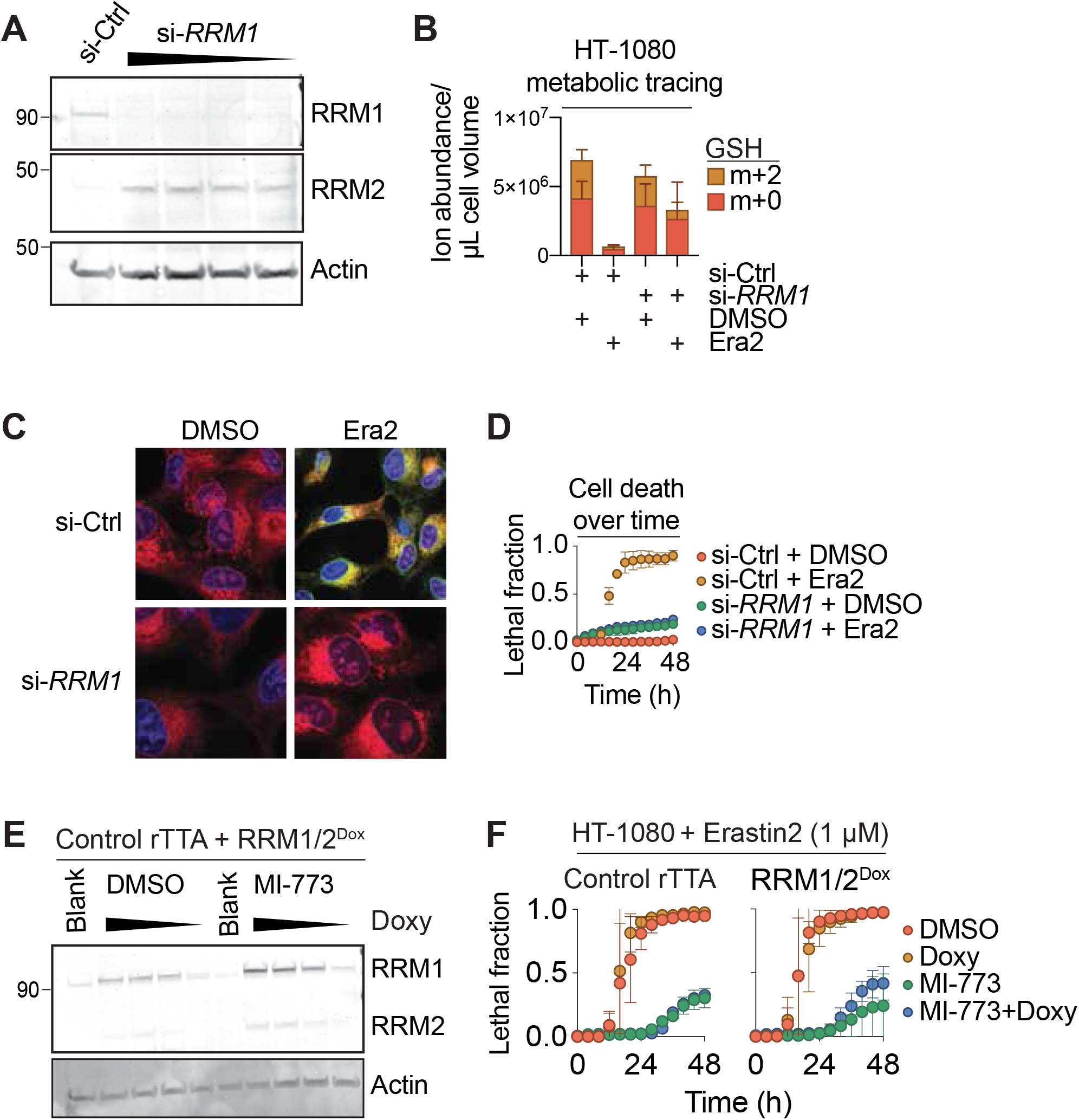
RNR expression modulates glutathione metabolism. **(A)** Protein expression in HT-1080 Control cells treated with non-targeting siRNAs (si-Ctrl), or an siRNA pool targeting *RRM1* (si-*RRM1*). Cells were pretreated with siRNA at doses ranging from 1 nM to 0.125 nM for 48 h prior to cell harvest. **(B)** Metabolic tracing of HT-1080 Control cells pretreated with si-Ctrl or si-*RRM1* for 48 h, then treated ± Era2 (1 μM, 8 h) in the presence of ^13^C-serine. De novo synthesized GSH is represented as an m+2 isotopologue (yellow). **(C)** Confocal C11-BODIPY 581/591 imaging of lipid peroxidation in HT-1080 Control cells pretreated with si-Ctrl or si-*RRM1* for 48 h, then treated ± Era2 (1 μM, 8 h) (1 μM, 9 h). Note: this timepoint was selected because it allowed for harvest of cells just prior to the onset of frank membrane permeabilization. **(D)** Cell death in HT-1080^N^ Control cells pretreated with -Ctrl or si-*RRM1* for 48 h, then treated ± Era2 (1 μM). **(E)** Protein expression in HT-1080^rTTA^ cells stably expressing doxycycline (Doxy)-inducible RRM1/2 (RRM1/2^Doxy^), pretreated with medium alone (Blank) or medium containing 10-fold dilutions of doxycycline (Doxy) from 1 μg/mL down to 1 ng/mL. **(F)** Cell death in HT-1080 cells pretreated ± nutlin-3 (10 μM) ± Doxy (100 ng/mL) for 48 h, then treated with erastin2. Data in **B, C** and **F** are mean ± SD of ≥3 biological replicates.

Next, we tested whether overexpression of RNR alone could restore normal ferroptosis sensitivity in cells where the p53/p21 pathway was activated. Towards this end, we generated a Dox-inducible overexpression construct driving the expression of *RRM1* and *RRM2* joined with a self-cleaving P2A cassette (HT-1080 RNR^Dox^). Dox treatment of HT-1080 RNR^Dox^ cells resulted in expression of both RRM1 and RRM2 at the expected sizes (**Figure 5E**). Optimization experiments suggested that nutlin-3 may weakly activate the expression of doxycycline-inducible cassettes non-specifically. Thus, in this experiment we employed the structurally distinct HDM2 inhibitor MI-733, which like nutlin-3 potently suppressed ferroptosis (**Figure 1C**). MI-733 pretreated cells were potently protected from erastin2-induced ferroptosis (**Figure 5F**). However, Dox-induced expression of RRM1 and RRM2 alone was not sufficient to revert this protective effect, likely because of coordinate downregulation of multiple nucleotide metabolic pathway genes in response to p53 stabilization (**Figure 3B**).

## Discussion

Nucleotide synthesis by RNR and related enzymes is a demanding task that consumes numerous metabolic resources. Of special note, the RNR reaction cycle consumes reducing equivalents supplied by glutathione (GSH), as well as thioredoxin, in mammalian cells (Sengupta *et al*., 2019; Zahedi Avval & Holmgren, 2009). GSH and thioredoxin also inhibit ferroptosis (Llabani *et al*, 2019; Yang *et al*., 2014). Thus, RNR-dependent nucleotide synthesis is in direct competition with GPX4-dependent lipid hydroperoxide reduction for use of the same co-substrates. It would therefore be expected that modulation of this competitive balance has the potential to shift cell sensitivity to ferroptosis. Indeed, our results suggest that inhibition of RNR function is sufficient to conserve intracellular GSH that can then be used to inhibit ferroptosis (**Figure 6**). Notably, proliferative arrest per se (e.g. using the CDK4/6 inhibitor palbociclib) is not sufficient to conserve GSH or inhibit ferroptosis (Tarangelo *et al*., 2018). Rather, it appears that direct inhibition of RNR (or direct upstream enzymes) is needed to accumulate ‘un-used’ GSH, which can be re-purposed by GPX4 to inhibit ferroptosis. How cells normally prioritize limited intracellular GSH pools for use in nucleotide synthesis versus lipid hydroperoxide reduction is an important direction for future investigation.

**Figure 6:**
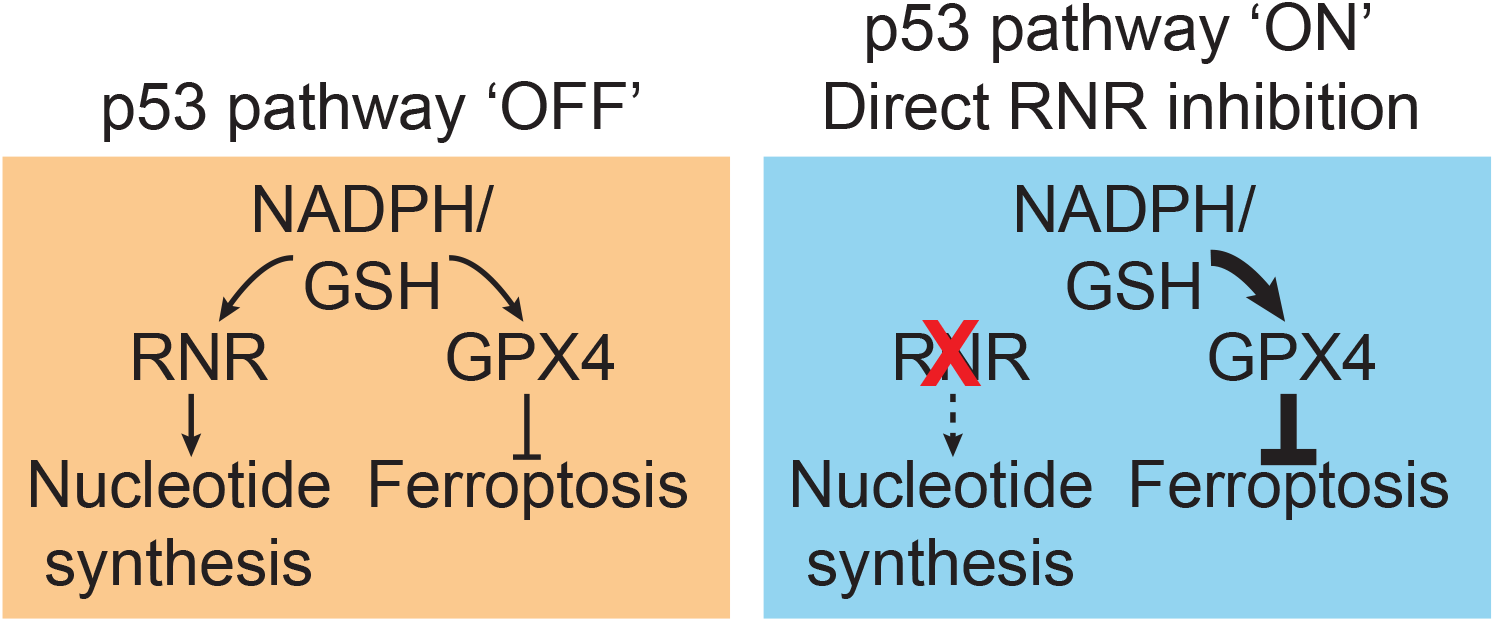
A model of p53-p21 activity in ferroptosis suppression. Stabilization of wild-type p53 leads to reduction of RNR expression and function, and conservation of intracellular GSH and potentially other reducing agents (e.g. NADPH), which can then be re-directed towards the inhibition of ferroptosis. Note: p53 may downregulate *RRM1* and *RRM2* in a partly p21-independent manner.

We propose that stabilization of wild-type p53 and induction of p21 can link downregulation of RNR-associated gene expression to protect against ferroptosis. Such a model would be consonant with, and elaborate upon, previous findings that wild-type p53 expression can protect against serine deprivation-associated oxidative stress (Maddocks *et al*., 2013). Of note, repression of RRM1/2 upon p53 stabilization was partly independent of p21. This may explain why p21 overexpression alone was not sufficient to fully inhibit ferroptosis in our hands: p53 stabilization is needed together with p21 induction for complete inhibition of RNR complex expression and function. However, others have observed that p21 expression alone can be sufficient to inhibit ferroptosis (Venkatesh *et al*., 2020b). It is possible that these differences are explained by cell type-specific differences, or the ability of p21 to regulate redox homeostasis via alternative means in some contexts (Chen *et al*, 2009). Regardless, these results point towards a meaningful role of p21 in regulating cell fate that is only partly dependent on p53.

In the context of cancer therapy, our results suggest that stimulation of nucleotide synthesis in combination with GSH depletion may be especially toxic. By contrast, the effects of anti-cancer agents that seek to induce ferroptosis via GSH depletion (e.g. cyst(e)inase, (Cramer *et al*, 2017)) could be blunted if combined with chemotherapeutics like gemcitabine and hydroxyurea that inhibit the consumption of GSH by nucleotide metabolic processes. Indeed, other conditions that slow intracellular metabolism, such as inhibition of mTOR signaling or deprivation of certain amino acids, also appear to cause ferroptosis resistance through GSH-conserving mechanisms that are unrelated to nucleotide synthesis (Conlon *et al*., 2021). Collectively, these studies suggest somewhat paradoxically that to maximize the induction of ferroptosis by GSH depletion it may be necessary to ensure that cancer cells are otherwise as metabolically active as possible.

## Author Contributions

Conceptualization, A.T., L.Z.L., and S.J.D. Methodology, A.T. and J.T.K.; Investigation, A.T., J.T.K.; Writing-Original Draft, A.T. and S.J.D.; Review & Editing, J.Z.L. Funding Acquisition and Supervision, S.J.D.

## Declaration of interests

S.J.D. is a member of the scientific advisory board of Ferro Therapeutics and holds patents related to ferroptosis.

## Acknowledgements

We thank Leslie Magtanong and Tim Stearns for help with confocal imaging, Jason Rodencal and Edo Biluar for reading the manuscript and assisting with experiments, and Julien Sage and Jan Carette for providing reagents. This work was supported by an NCI F99/K00 Predoctoral to Postdoctoral Transition Award to A.T. (1F99CA234650-01) and an award from the NIH to S.J.D. (1R01GM122923).

## Experimental Procedures

### Cell lines

HT-1080 (CCL-121), U-2 OS (HTB-96), and 293T (CRL-3216) were obtained from ATCC (Manassas, VA, USA). Cell viability studies used human polyclonal cell lines expressing nuclear-localized mKate2 (Nuc::mKate2). Nuc::mKate2 cell lines were generated by transducing parental cell lines with NucLight Red lentivirus (Essen Biosciences, Ann Arbor, MI, USA). Polyclonal Nuc::mKate2-expressing HT-1080 cells (denoted HT-1080^N^), as well as similar U-2 OS^N^ cell lines, were described previously (Forcina *et al*., 2017). Transduced cell lines were selected with 1 μg/mL puromycin (Cat# A11138-03, Fisher Scientific, Hampton, NH) for 3 days. HT-1080 *TP53* and *CDKN1A* knockout (KO) cell lines generated using CRISPR-Cas9 genome editing were described previously (Tarangelo *et al*., 2018). Rho zero (ρ^0^) HT-1080 cells were generated as previously described (Dixon *et al*., 2012). Briefly, HT-1080 cells were cultured in media containing 10 ng/mL ethidium bromide and 50 μg/mL uridine for 14 passages. Uridine was maintained in the media for all experiments and was also supplemented into the medium of control ρ^+^ cells.

### Cell culture conditions

HT-1080 and 293T cells were cultured in DMEM Hi-glucose medium (Cat# MT-10-013-CV, Corning Life Science) supplemented with 1% non-essential amino acids (NEAAs) (Cat# 11140-050, Life Technologies, Carlsbad, CA, USA). U-2 OS cells were cultured in McCoy’s 5A media (Cat# MT-10-050-CV, Corning Life Science, Tewksbury, MA, USA). All media was supplemented with 1x Pen/Strep (Life Technologies) and 10% fetal bovine serum and filtered through a 0.22 μM PES filter (Genesee Scientific, San Diego, CA, USA). All cell lines were grown at 37°C with 5% CO_2_ in humidified tissue culture incubators (ThermoFisher Scientific, Waltham, MA, USA).

### Chemicals and reagents

Erastin2 and ML162 were synthesized at Acme Bioscience (Palo Alto, CA, USA). Nutlin-3 (Cat# S1061), Nutlin-3b (Cat# S8065), gemcitabine HCL (Cat# S1149) and MI-773 (Cat# S7649) were purchased from Selleck Chemicals (Houston, TX, USA). Pyrazofurin (Cat# SML1502), hydroxyurea (Cat# H8627), mycophenolic acid (Cat# M5255), ferrostatin-1 (Cat# SML0583), doxycycline hyclate (Cat# D9891), and L-buthionine sulfoximine (Cat# B2515) were purchased from Sigma-Aldrich. Buthionine sulfoximine and gemcitabine HCL were dissolved directly into cell media. C11 BODIPY 581/591 was prepared as a stock solution in methanol and stored prior to use at −20°C. All other compounds were prepared as stock solutions in DMSO and stored prior to use at −20°C.

### Cell viability experiments

All experiments examining the effects of p53 activation on cell death utilized a pretreatment phase (24 or 48 h). On day one, cells were seeded into clear-bottom 96-well assay plates (Cat# 07-200-588, Fisher Scientific). Cell seeding was optimized to achieve approximately equal densities of cells in each well, accounting for the cytostatic effects of some compounds during the pretreatment phase. The following number of cells per well were seeded. HT-1080 Control^N^: DMSO, 1000-1500; Nutlin-3 family treatment, 3000-4000; gemcitabine, 3000-4000). HT-1080^N^ p53^KO^: DMSO, 750-1000; Nutlin-3 family treatment, 750-1000; gemcitabine, 2000, HT-1080^N^ p21^KO^: DMSO, 2000-3000; Nutlin-3 family treatment, 4000-5000; gemcitabine, 4000. The pretreatment phase began on day 2 when media was replaced with media containing DMSO or a small molecule inhibitor (e.g. nutlin-3, 10 μM or gemcitabine HCL, 200 nM). During the pretreatment phase, cell death was not monitored. On day 4, following 48 h of pretreatment, media was removed and replaced with media containing lethal compounds or other modulating compounds (such as buthionine sulfoximine, 100 μM). DMSO or pretreatment compounds were maintained in the medium during the observed treatment phase. During the treatment phase, cell death was observed and quantified using scalable time-lapse analysis of cell death kinetics (STACK) as previously described (Forcina *et al*., 2017).

### Cell death kinetics quantification

Analysis of cell death kinetics was performed using STACK analysis (Forcina *et al*., 2017). Following pretreatment, infection, or transfection, cell media was exchanged for media containing lethal compounds and 0.022 μM of the viability dye SYTOX Green (SG, Cat # S7020, Life Technologies). Cells were then imaged at 4 h intervals for ≥ 48 h using the Essen IncuCyte ZOOM live cell analysis system (Essen BioSciences). Live cells were quantified in HT-1080^N^ or U-2 OS^N^ cells by counting mKate2 positive cells, and dead cells were quantified by counting SG positive cells. All quantification was performed automatically using the IncuCyte ZOOM Live-Cell Analysis System software (Essen BioSciences). Lethal fraction scores were calculated at each time point as previously described, with two modifications (Forcina *et al*., 2017): (*i*) Because long-dead cells can release SG, the maximum number of SG positive counts from the start of the time course was used in these calculations; (*ii*) SG/mKate2 double-positive cells were considered to be in the process of death, and thus counts of double positive cells were subtracted from the counts of live cells. Lag exponential death (LED) curves were fitted to lethal fraction data using Prism 8 (GraphPad, La Jolla, CA, USA), as described (Forcina *et al*., 2017). The parameter of time to death onset, or D_O_ was computed from LED curve fits, as previously described (Forcina *et al*., 2017).

### Immunoblotting

Cells seeded in 6-well dishes were washed twice with HBSS (Cat# 14025-134, Life Technologies) and collected using a manual cell lifter. Cell pellets were collected and lysed in 9M urea or RIPA buffer (10 mM Tris ph7.5, 150 mM NaCl, 1 mM EDTA, 0.5% Na Deoxycholate, 0.1 % SDS, 1% Triton X). Lysates were sonicated 10 times with one-second pulses at 80% amplitude on a Fisher Scientific Model 120 Sonic Dismembrator (Thermo Fisher), then centrifuged at ≥13,000 RPM at room temperature (for 9M urea) or at 4°C (for RIPA lysis) for 15-20 min. Lysates were quantified by BCA assay (Cat # PI23224, Thermo Scientific). Samples were prepared with NuPage 10x Reducing Agent (Cat# NP0009, Thermo Fisher) and NuPage LDS 4x Sample Buffer (Cat# NP0007, Thermo Fisher). Samples were incubated at 70°C for 10 minutes and run on pre-cast NuPage SDS 4-12% gradient gels in NuPage MES Running Buffer (Cat# NP0002, Thermo Fisher). Gels were transferred to nitrocellulose membranes using an iBlot Dry Blotting System (Cat# IB21001, Thermo Fisher), blocked for ≥1 h in 50% Odyssey PBS (Cat# 927-40010, Li-Cor) /50% diH_2_O, and probed with primary antibodies either overnight at 4°C or at room temperature or 1-3 h. Primary antibodies used were against actin (Cat# SC-1616, I-19 or Cat # sc-47778, C4, Santa Cruz, 1:1000), human p53 (Cat #SC-126, DO-1, Santa Cruz, 1:1000), p21 C’-terminus (Cat# 2947, 12D1, Cell Signaling Technology, 1:1000), p21 N’-terminus (Cat # ab80633, CP74, Abcam, 1:500), RRM1 (Cat # CST3388, Cell Signaling Technologies, 1:500), and RRM2 (Cat # 65939, Cell Signaling Technologies, 1:500). Membranes were washed three times for 10 min each in Tris buffered saline (Cat# 0788, ISC BioExpress) with 0.1% Tween 20 (TBST) at room temperature with rocking. Membranes were probed with secondary antibodies (Donkey anti-goat-680 Cat# 926-68024, Donkey anti-goat-800 Cat# 926-32214, Donkey anti-rabbit-680 Cat# 926-68023, Donkey anti-rabbit-800 Cat# 926-32213, Donkey anti-mouse-680 Cat# 926-68022, Donkey anti-mouse-800 Cat# 926-32212, Li-Cor) in 50% Odyssey Buffer (Li-COR, Cat# 927-40100)/50% diH_2_O with 0.1% SDS and 0.4% Tween 20 for 1 h at room temperature with rocking. Membranes were washed three times for 10 minutes each in TBST, then imaged on a Li-COR Odyssey CLx imager.

### Inducible gene expression and cloning

HT-1080 cell lines expressing tetracycline inducible full-length or truncated forms of p21^20^ were generated as follows. gBlocks containing a FLAG tag followed by the full coding sequence of human *CDKN1A*, the N’-terminus only (amino acids 1-90), or the C’-terminus (amino acids 87-164) were synthesized by IDT DNA (Coralville, IA) and cloned into lentiviral vectors containing a Tet-inducible promoter (pLenti-CMV-TRE3G-Puro, a gift of Jan Carette) by Gibson assembly. HT-1080^N^ cells were transduced with lentiviruses carrying the reverse tetracycline-controlled transactivator 3 under a CMV promoter (CMV-rtTA3) (Cat# w756-1, Addgene). HT-1080^N-rtTA3^ cell lines were selected in 10 μg/mL blasticidin (Cat # A1113902, Thermo Fisher Scientific) for 5 days. These cell lines were then transduced with lentiviruses carrying full length or truncated FLAG-tagged p21 coding sequences under a Tet-inducible promoter. Cell lines were selected with 1 μg/mL puromycin for 3 days.

To overexpress ribonucleotide reductase, the following construct was synthesized as gBlocks and assembled using Gibson assembly (Master Mix Cat # E2611S, New England Biolabs, Ipswich, MA, USA). The FLAG-tagged coding sequence of *RRM1* was joined via a P2A cassette to the 6x-His tagged coding sequence of *RRM2*. This construct was cloned into the pLenti-CMV-TRE3G vector (pLenti-CMV-TRE3G-FLAG-RNR-P2A-HIS-RRM2). HT-1080^N-rtTA3^ cell lines were infected with viruses carrying pLenti-CMV-TRE3G-FLAG-RNR-P2A-HIS-RRM2 and selected using 1 μg/mL puromycin for 3 days.

To induce gene expression in tetracycline-inducible cell lines, cells were treated with 1 ng-1 μg doxycycline for 24-48 h. Gene expression was confirmed using immunoblotting as described above.

### Lentivirus production

To generate lentiviruses, 293T cells were seeded at a density of 0.5 x 10^6^ cells in a 6-well dish the day before transfection. Cells were transduced the next day with 1000 ng of plasmid DNA plus 250 ng pMD2.G (Cat# 12259, Addgene) and 750 ng psPax2 (Cat# 12260, Addgene) 2^nd^ generation lentiviral packaging plasmids. Transfection was performed using 3 μL PolyJet (Cat# SL100688, SignaGen Laboratories, Rockville, MD, USA) transfection reagent diluted in plain DMEM to a final volume of 100 μL and incubated for 15 minutes at room temperature. The transfection mixture was added drop-by-drop to wells and incubated for 24 h. After 24 h, the cell supernatant was removed and replaced with 1.5-2 mL fresh complete DMEM. After 8 h, virus-containing media was harvested and replaced with 1.5-2 mL fresh complete DMEM. The following day, cell media was harvested twice more at an interval of >8 h, for a total of three harvests. Virus-containing media was filtered using a 0.45 μM PVDF (Cat# SLHV033RS, Millex, Duluth, GA, USA) syringe-driven filter unit and frozen at −80°C until use. Relative viral titer was determined by infecting cells with serially diluted volumes of virus-containing media and selecting using appropriate selection reagents.

### siRNA gene knockdown

Cells were reverse-transfected with an ON-TARGET Plus siRNA SMARTPool targeting human *RRM1* (targeting NM_001033.5, NM_001318064.1, NM_001318065.1, NM_001330193.1, Cat # L-004270-00-0005, Horizon, Summerville, SC, USA). AllStars Negative Control siRNA (Cat# 1027280, Qiagen, Redwood City, CA, USA) was used as a negative control. AllStars Hs Cell Death siRNA (Cat# 1027298, Qiagen) was used as a positive control for transfection efficiency. Transfection mixes were prepared using 0.25 nM siRNA and 2.5 μL Lipofectamine RNAiMAX Transfection Reagent (Cat# 13778075, Life Technologies) to a total volume of 500 μL transfection mixture in Opti-MEM Reduced Serum Media (Cat# 31985-062, Life Technologies). 250 μL Opti-MEM was pipetted into empty 6 well dishes and siRNA was added directly to the wells. 250 μL of lipofectamine mixture was added to the siRNA mixture. The combined mixture was swirled and incubated 15 minutes at room temperature. HT-1080 cells were seeded in a 6-well at the following densities per well. HT-1080 Control: siControl, 0.5-1 x 10^5^; si*RRM1*, 1.5-2 x 10^5^. HT-1080 p53^KO^ and p21^KO^ cell lines were seeded in a 12-well dish at the following densities. HT-1080 p53^KO^: siControl, 0.15 x 10^5^; si*RRM1*, 0.7 x 10^5^. HT-1080 p21^KO^: siControl, 0.3 x 10^5^; si*RRM1*, 1 x 10^5^. HT-1080 cells were seeded in a 6-well at the following densities per well: siControl, 0.6 x 10^5^; si*RRM1*, 1.5 x 10^5^. Cells were added in a volume of 1.5 mL, for a total well volume of 2 mL. Cells were incubated for 48 h prior to assay.

### C11 BODIPY 581/591 imaging

HT-1080 cells were seeded on glass coverslips in 6-well dishes at the following densities: Blank Media, 0.3 x 10^5^; Gemcitabine, 1 x 10^5^; siControl, 0.5 x 10^5^; si*RRM1*, 2 x 10^5^. For the *RRM1* knockdown condition, reverse transfection reagent was prepared on top of glass cover slips in 6-well dishes using either non-targeting control siRNAs, or siRNA pools targeting human *RRM1*, as described above. On day 2, media was removed and replaced with fresh complete media. For the Gemcitabine treatment condition, cells were seeded on Day 1 as described above. On Day 2, cells were pretreated with or without Gemcitabine HCL (200 nM). For both conditions, on day 4, cell mediate was removed and replaced with media containing DMSO or 1 μM Erastin2. Cells were incubated at 37° for 9.5 hours, prior to the onset of cell death, after which point cell media was removed and cells were washed with HBSS. C11-BODIPY 581/591 was prepared as a 5 μM working solution in HBSS and added to cells along with 1 μg/mL Hoechst 33258 stain (Cat # H3569, Thermo Fisher). Cells were incubated for 10 min at 37°C, then the C11-BODIPY mixture was removed and cells were washed in 1 mL of fresh HBSS. To prepare slides for imaging, 25 μL HBSS was pipetted onto a microscope lined in parafilm. Coverslips with stained cells were lifted out of plates using a needle tip and inverted onto prepared slides. Slides were sealed with melted Vaseline and immediately imaged on a Zeiss Observer Z1 confocal microscope. Images were processed in ImageJ. Brightness for the green (oxidized BODIPY) channel was auto-scaled using images from cells treated with erastin2. Brightness for all images was auto-adjusted based on imaged with the brightest signal (i.e. brightness for the red (reduced BODIPY) channel was auto-scaled using images from cells treated with DMSO, while brightness for the green (oxidized BODIPY) channel was auto-scaled using images of cells treated with erastin2).

### Image analysis

Image processing was performed in Image J (version 1.50i) or Adobe Photoshop CS6 (Adobe Systems, San Jose, CA).

### Reverse transcription and quantitative PCR

Cells were seeded in 6-well dishes and treated with desired reagents. Following treatment, cells were washed twice with HBSS and collected using a cell lifter. Pellets were centrifuged at room temperature for 2 min at 3000 RPM. Supernatants were removed and lysed for RNA extraction using a Qiashredder extraction column (Qiagen, Cat# 79654) and the RNeasy Plus RNA Extraction Kit (Cat# 74134, Qiagen). Reverse transcription to generate cDNA was performed using TaqMan Reverse Transcriptase Kit according to the manufacturer’s instructions (Cat# N8080234, TaqMan). Quantitative polymerase chain reactions (PCR) were prepared using SYBR Green Master Mix (Cat# 4367659, Life Technologies) and run on an Applied Biosystems QuantStudio 3 real-time PCR machine (Thermo Fisher). Relative transcript levels were calculated using the ΔΔCT method and normalized to the *ACTB* gene. Primer sequences are as follows: *ACTB*, *forward*: ATCCGCCGCCCGTCCACA, reverse: ACCATCACGCCCTGGTGCCT; *CKDN1A*, forward: CACCGAGACACCACTGGAGG, reverse: GAGAAGATCAGCCGGCGTTT; *MDM2*, forward: GAATCATCGGACTCAGGTACATC, reverse: TCTGTCTCACTAATTGCTCTCCT; *RRM1*, forward: CAGTGATGTGATGGAAGA, reverse: CTCGGTCATAGATAATAGCA; *RRM2*, forward: AGACTTATGCTGGAACT, reverse: TCTGATACTCGCCTACTC; *ND1* forward: CCACATCTACCATCACCCTC, reverse: TTCATAGTAGAAGAGCGATGGT; *ND2, forward:* GACATCCGGCCTGCTTCTT, reverse: TACGTTTAGTGAGGGAGAGATTTGG; *ND5*, forward: TTCAAACTAGACTACTTCTCCATAATATTCATC, reverse: TTGGGTCTGAGTTTATATATCACAGTGA.

### RNA sequencing

HT-1080 cells were seeded in 6-well dishes at the following densities. Control HT-1080s: DMSO, 0.5 x10^5^, Nutlin-3, 1.5 x10^5^. HT-1080 p53^KO^: DMSO, 0.3 x10^5^, Nutlin-3, x10^5^. HT-1080 p21 ^KO^: DMSO, 0.6 x10^5^, Nutlin-3, 1.5 x10^5^. Cells were treated for 48 hr with DMSO or 10 μM nutlin-3. Cells were then washed with ice-cold HBSS and detached with a cell lifter on ice. Detached cells were pelleted by centrifugation at 3000 RPM for 2 min. The supernatant was removed and cell pellets were immediately lysed and RNA was extracted using a (Qiagen, Cat# 79654) and the RNeasy Plus RNA Extraction Kit (Cat# 74134, Qiagen). Two biological replicates were collected for each sample. Purity, concentration, and integrity of RNA was assessed by NanoDrop Spectrophotometer (Thermo Fisher Scientific) and by Eukaryote Total RNA Nano chip analysis on a Bioanalyzer (Agilent) at the Stanford Protein and Nucleic Acid Facility. The RNA Integrity Number for each sample met or exceeded the minimum threshold of 6.8, and NanoDrop 260/280 and 260/230 ratios achieved a threshold of at least 1.95. Following quality control steps, samples were shipped on dry ice to Novogene (Sacramento, CA) for library generation and 20M read PE150 sequencing on an Illumina HiSeq 4000 Platform. Data cleanup was performed by excluding reads with: adaptor contamination; >10% of indeterminate bases, or >50% bases with a quality score ≤ 5. The remaining reads (≥98% of all reads across all conditions) were aligned to the hg19 human reference genome using STAR. At least 91% of clean reads were mapped across all conditions. Pearson correlation between biological replicates was R^2^ ≥ 0.96 for all samples. Differential gene expression was calculated using DESeq 1.10.1. We selected all protein-coding genes with read counts above 0.5 FPKM in all conditions and with significant alterations (adjusted P value < 0.05).

### Metabolic tracing analysis

For these experiments, cells were cultured in RPMI 1640 medium (Cat# SH30027FS, Fisher Scientific) containing 10% FBS and 1x Pen/Strep. HT-1080^N^ cells were seeded in 6-well dishes at the following densities per well. Control HT-1080^N^: DMSO, 0.3 x 10^5^, gemcitabine, 1.5-2 x 10^5^. HT-1080^N^ p53^KO^: DMSO, 0.25 x 10^5^, gemcitabine, 0.75 x 10^5^. HT-1080^N^ p21^KO^: DMSO, 0.4 x 10^5^, gemcitabine, 1.5 x 10^5^. The following day, cells were pretreated with blank media or gemcitabine HCL (200 nM) for 48 h in RPMI 1640 medium. For iterations of this experiment ± Gemcitabine (200 nM) conditions were maintained from the pretreatment phase. Cells were incubated in [U-^13^C] L-serine media for 8 h. Unlabeled controls were incubated for 8 h in standard RPMI 1640 medium. In a separate iteration of this experiment, HT-1080 cells were reverse transfected with non-targeting (control) siRNAs or siRNAs targeting *RRM1* and seeded as described above. After 48 h, media was removed and replaced with RPMI medium lacking glucose, serine, and glycine (Cat# R9660-02, TEKnova) supplemented with 30 mg/L [U-^13^C] L-Serine (Cat# 604887, Sigma-Aldrich) and 2000 mg/L D-glucose (Cat# G54000, Sigma-Aldrich) ± erastin2 (1 μM). Prior to collection, cells were imaged on the IncuCyte ZOOM and total cell number of the well was computed based on counts of the total mKate2^+^ objects per area of viewing field. After the treatment phase, cells were washed twice in ice cold HBSS, and fixed in 80% cold LC/MS grade acetonitrile (Cat #A955-500, Fisher Scientific) for 5 min on ice. Cells were collected using a cell lifter, centrifuged at ≥12,000 RPM for 10 min at 4°, and the resulting supernatant was removed to glass vials and stored at −80°C prior to analysis. The remaining pellets were frozen at −80°. Due to the effect of gemcitabine and *RRM1* knockdown on cell size, ion abundance was normalized to total cell volumes. To quantify cell volume, cells were seeded and treated exactly as described above. After appropriate pretreatment phases, cells were trypsinized, centrifuged for 5 min at 1,500 RPM, and resuspended in 500 μL ISOTON Diluent (Cat# 12754878, Fisher Scientific). Cell suspensions were diluted 250 μL into 10 mL of ISOTON diluent and cell size was measured on a Coulter Z2 Cell Analyzer (Cat# 844 80 11, Beckman, Indianapolis, IN). Median cell volume was used for calculations. Median cell volume was multiplied by the number of cells counted prior to collection to yield a total cell volume for each sample. Ion abundances were normalized to cell volumes.

For separation of polar metabolites, normal-phase chromatography was performed with a Luna 5 μm NH2 100 Å LC column (Phenomenex 00B-4378-E0). Mobile phases were as follows: Buffer A, acetonitrile; Buffer B, 95:5 water/acetonitrile with 0.2% ammonium hydroxide and 50mM ammonium acetate for negative ionization mode. The flow rate for each run started 100% B for 2 minutes at 0.2 mL/min, followed by a gradient starting at 20% A / 80% B changing linearly to 70% A / 30% B over the course of 1 minute at 0.7 mL/min, followed by 70% A / 30% B for 5 minutes at 0.7 mL/min and finally to 100% B over a course of 5 minutes. The overall runtime was 13 min and the injection volume was 20 μL.

MS analysis was performed with an electrospray ionization (ESI) source on an Agilent 6545 qTOF LC/MS at the Metabolic Chemistry Analysis Center at Stanford ChEM-H. For QTOF acquisition parameters, the mass range was set from 50 to 1000 m/z with an acquisition rate of 5 spectra/ second and time of 250 ms/ spectrum. For Dual AJS ESI source parameters, the drying gas temperature was set to 250°C with a flow rate of 12 l/min, and the nebulizer pressure was 20 psi. The sheath gas temperature was set to 300°C with a flow rate of 12 l/min. The capillary voltage was set to 3500 V and the fragmentor voltage was set to 100 V. Isotopologues extraction was performed in Agilent Profinder B.08.00 (Agilent technologies). Retention time (RT) of each metabolite was determined by authentic standards (Table). The mass tolerance was set to +/-15 ppm and RT tolerance was +/-0.2 min. Glutathione (formula C10H17N3O6S, METLIN: 44, HMP: HMDB00125, KEGG C00051) was detected with a mass of 307.084 and a retention time of 8.78 minutes.

### Seahorse assay

HT-1080 cells were seeded in Seahorse XFp 6-well miniplates at the following densities. HT-1080 Control: DMSO, 750 cells; Nutlin-3, 2000 cells; Palbociclib, 1500 cells. HT-1080 p53^KO^: DMSO, 500 cells; Nutlin-3, 500 cells; Palbociclib, 500 cells. HT-1080 p21^KO^: DMSO, 1000 cells; Nutlin-3, 2000 cells; Palbociclib, 1500 cells. Cells were pretreated with small molecules in Seahorse XFp plates for 48 h prior to assay. For ρ^0^ validation, 8000 HT-1080 Control ρ^+^ and 12,000 HT-1080 Control ρ^0^ cells were seeded into XFp 6-well mini plates and assayed the next day. XFp Mitochondrial and Glycolytic Stress Tests were performed per the manufacturer’s instructions. All data was normalized to cell number as determined by counts of NLR^+^ cells prior to the start of the assay.

### Graphing and figure assembly

Graphing and statistical analyses were performed using GraphPad Prism 8. The Results and individual Figure Legends contain additional statistical details. Figures were constructed in Adobe Illustrator.

### Data analysis

Data analysis was performed using Microsoft Excel and GraphPad Prism 8. Data are represented as mean ± SD or ± 95% confidence interval unless otherwise noted.

---

**Supplemental Figure 1.**
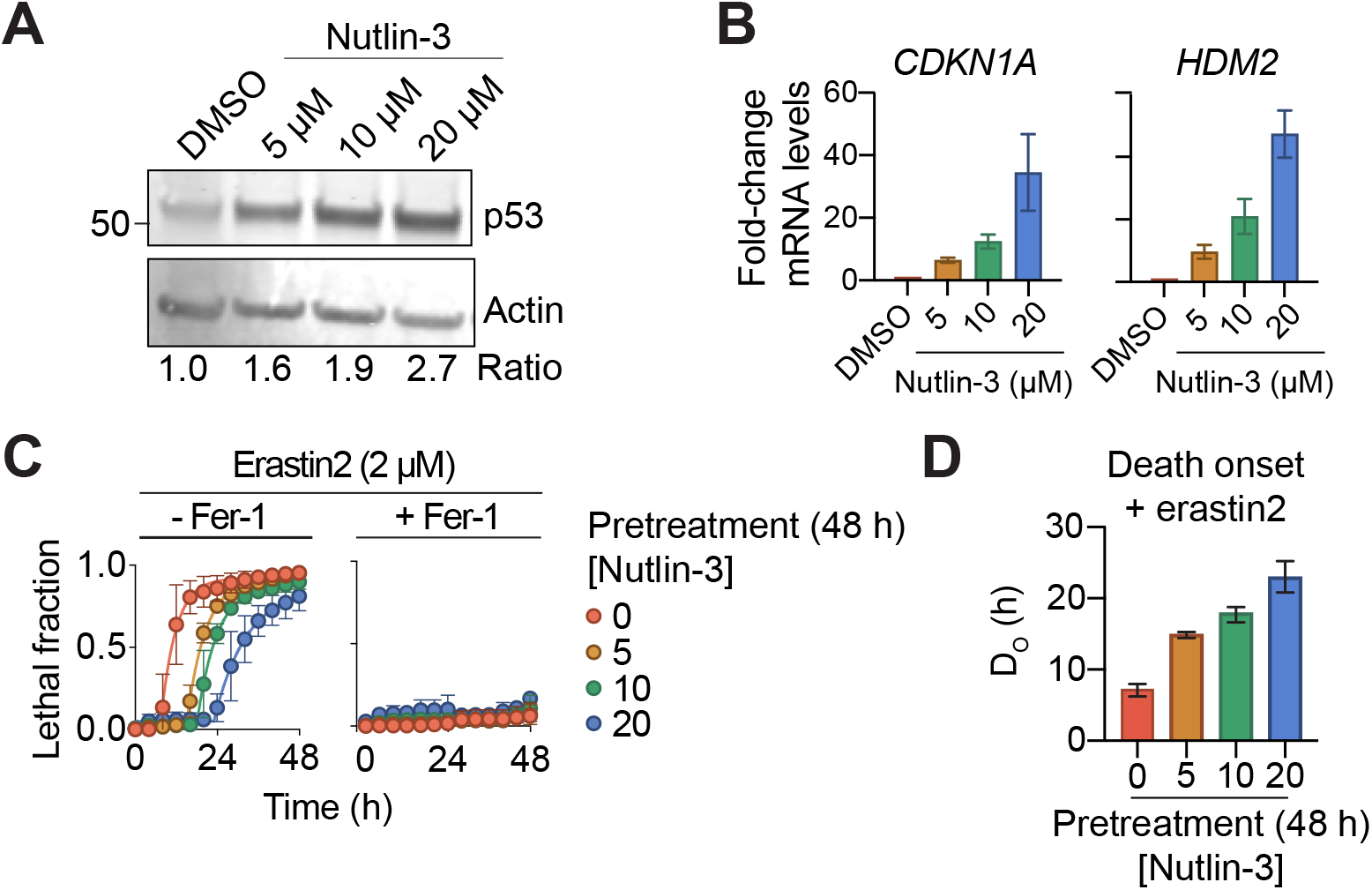
p53 stabilization suppresses ferroptosis. (A) Protein expression in HT-1080 cells in response to nutlin-3 treatment (48 h). (B) Relative mRNA expression determined using RT-qPCR in HT-1080 cells. (C) Cell death over time ± erastin2 ± ferrosta-tin-1 (Fer-1, 1 μM) following pretreatment with increasing concentrations of nutlin-3. (D) Timing of cell death onset (D_O_), determined using STACK for the lethal fraction curves in C. Results are mean ± SD (B,C) or mean ± 95% C.I. from three independent experiments.

**Supplemental Figure 2.**
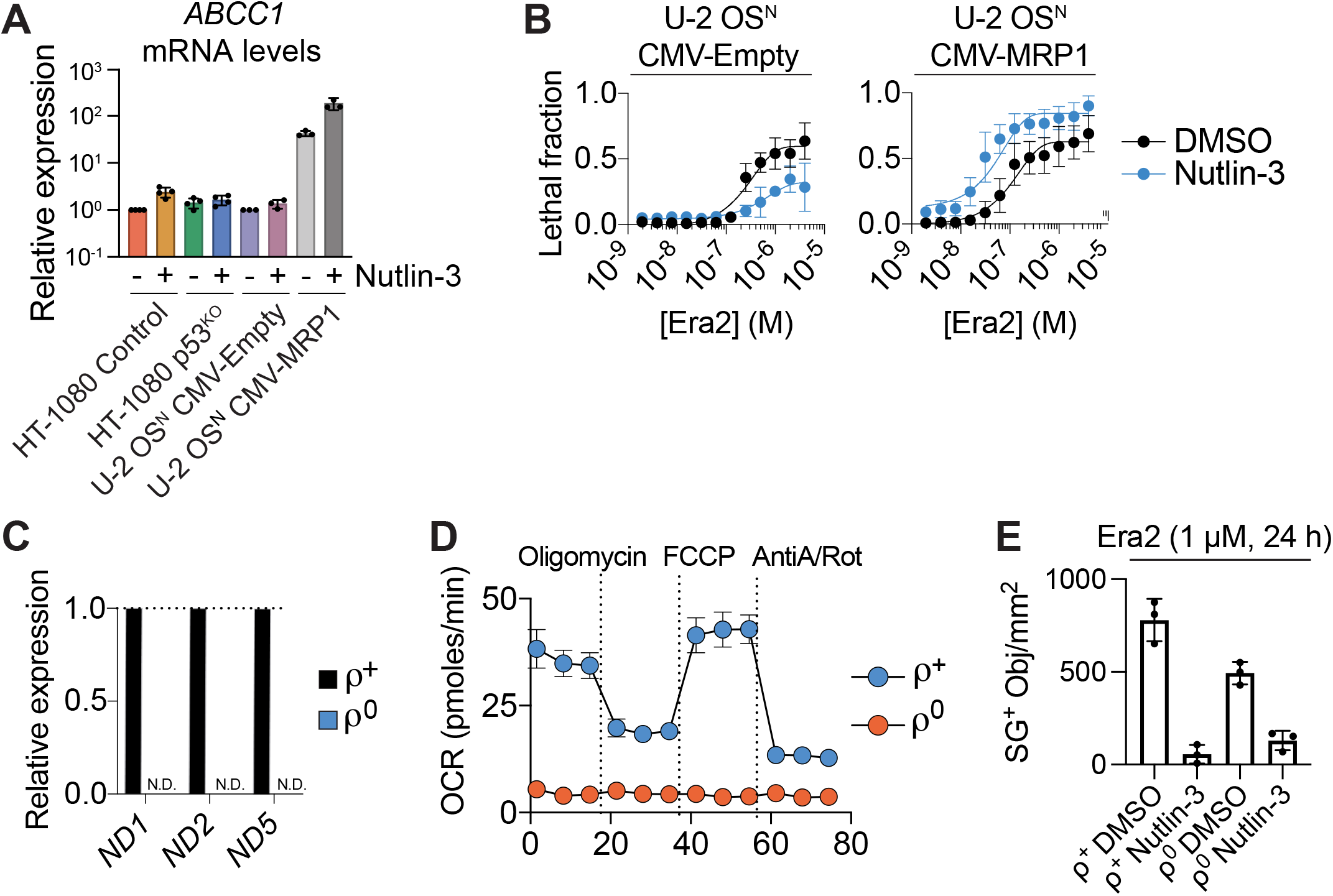
p53 stabilization does not suppress ferroptosis via effects on glutathione export or mitochondrial function. (A) Relative mRNA expression determined using RT-qPCR. Nutlin-3 was used at 10 μM (48 h). (B) Cell death in cells that overexpress an empty control vector (CMV-Empty) or MRP1 (ABCC1) (CMV-MRP1), following pretreatment ± nutlin-3 (10 μM, 48 h) then treatment with erastin2 (Era2, 48 h). (C) Relative mRNA expression for three mitochondrial DNA-encoded transcripts in HT-1080 rho (ρ) positive or rho negative cell lines. (D) Oxygen consumption rate (OCR) determined in HT-1080 ρ^+^ and ρ^0^ cell lines, as determined using Seahorse technology. AntiA: antimycin A, Rot: rotenone. (E) SYTOX Green positive (SG^+^) dead cell counts in HT-1080 ρ^+^ and ρ^0^ cell lines pretreated ± nutlin-3 (10 μM, 48 h), then treated with Erastin2 (Era2). Data are mean ± SD (A-D) or mean ± 95% C.I. (E) from three independent experiments.

**Supplemental Figure 3.**
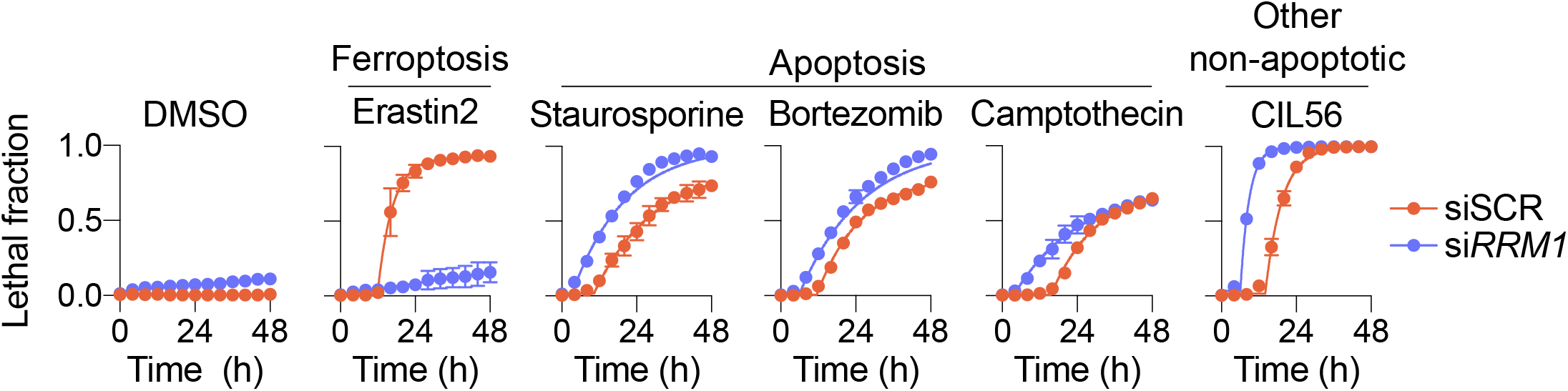
*RRM1* silencing specifically impacts ferroptosis. HT-1080 cells were treated with a scrambled control siRNA (siSCR) or an siRNA targetting RRM1 (si*RRM1*) for 48 h, then cell death was monitored in response to the indicated compounds. Lethal compounds were used at the following concentrations: erastin2 (1 μM), staurosporine (1 μM), bortezomib (0.2 μM), campthothecin (3 μM), CIL56 (10 μM). Data represents mean ± SD from three independent experiments.

